# GCase deficiency impairs uptake of α-synuclein-loaded ectosome by astrocytes

**DOI:** 10.64898/2026.11.15.766708

**Authors:** Julie Jacquemyn, Diana U. Rodrigues, Ehlam Iftikhar, Jinlan Chang, Vincent Soubannier, Thomas M. Durcan, Maria S. Ioannou

## Abstract

Parkinson’s disease (PD) involves the prion-like propagation of α-synuclein. One mechanism for the spread of α-synuclein between neurons in PD is ectosomes. But PD is a multicellular disorder in which astrocytes also play a critical role. How astrocytes take up ectosomes and whether this is affected in PD is unclear. Here we show that GCase-deficiency, as occurs in PD, impairs astrocytic uptake of neuron-derived ectosomes containing α-synuclein. The defect in ectosome uptake by astrocytes is driven by compromised macropinocytosis and can be rescued by reducing glucosylceramides or restoring lysosomal function. These findings identify sphingolipid metabolism as a key regulator of ectosome uptake by primary and iPSC-derived astrocytes. They also support a shift from the uptake and clearance of α-synuclein by astrocytes in health, towards the transmission of α-synuclein pathology towards neurons in PD.

**GRAPHICAL ABSTRACT:** 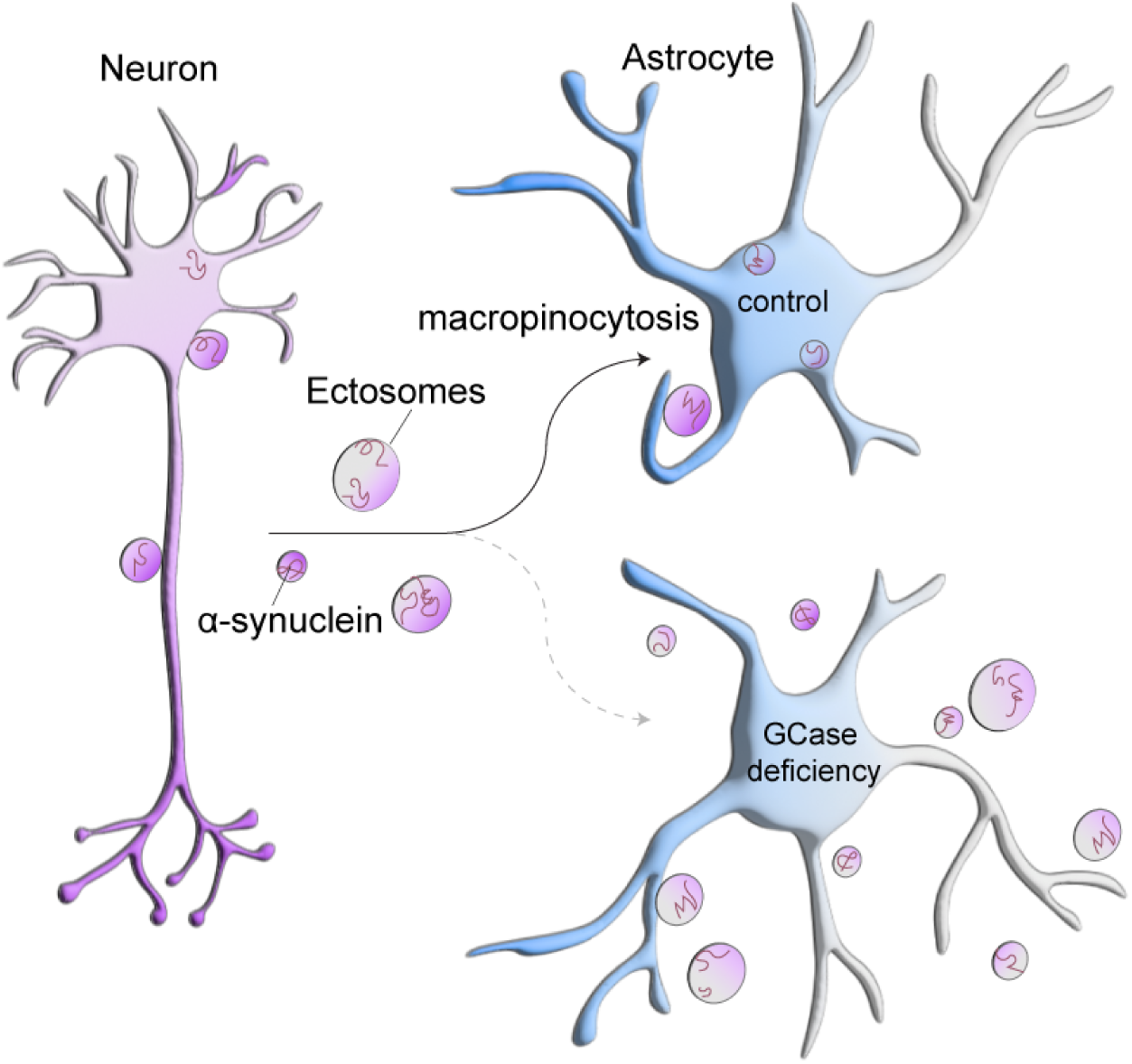

**HIGHLIGHTS:**

- GCase-deficiency impairs astrocytic but not neuronal uptake of lipids
- GCase-deficient astrocytes have impaired uptake of ectosomes containing α-synuclein
- Impaired uptake by astrocytes mediated by reduced macropinocytosis
- Reducing glucosylceramide or restoring lysosomal function rescues ectosome uptake

## INTRODUCTION

Parkinson’s disease (PD) is a progressive neurodegenerative disorder characterized by the selective degeneration of dopaminergic neurons and the intracellular accumulation of misfolded α-synuclein in Lewy bodies and Lewy neurites^1–3^. Although historically considered a neuron-centric disorder, PD is now recognized as a multicellular disease in which non-neuronal cells such as astrocytes, play critical roles in disease initiation and progression^4,5^.

A substantial body of evidence supports a prion-like mechanism underlying PD pathogenesis, whereby misfolded α-synuclein propagates between cells, seeding aggregation in recipient cells^3,6,7^. This cell-to-cell transmission is increasingly viewed as a central driver of disease progression and correlates with the stereotyped spatiotemporal spread of pathology observed in the human brain^1,2^. However, the cellular and molecular mechanisms governing α-synuclein release, transfer, and uptake and the relative contribution of the different brain cell types remain incompletely defined.

In parallel, dysregulated lipid metabolism has emerged as a fundamental feature of PD^8–11^, with astrocytes occupying a central role in brain lipid metabolism. Notably, mutations in *GBA1*, encoding the lysosomal hydrolase β-glucocerebrosidase (GCase), represent the strongest genetic risk factor for PD^12–14^. Loss of GCase function results in reduced enzymatic activity, accumulation of its substrate glucosylceramide, and widespread perturbations in lipid metabolism^15–25^. Reduced GCase activity and elevated glucosylceramide levels are not restricted to *GBA1*-associated PD but are also observed in *LRRK2*-associated and sporadic PD^26,27^, indicating that sphingolipid dysregulation constitutes a convergent and potentially causal mechanism in PD pathogenesis. Importantly, astrocytes are highly sensitive to disruption in sphingolipid homeostasis^28–30^.

Amongst other pathways, extracellular vesicles (EVs) have been proposed as mediators of α-synuclein intercellular transfer^31^. We recently demonstrated that reduced GCase activity in PD stimulates shedding of plasma membrane derived vesicles, ectosomes. These neuron-derived ectosomes are enriched in misfolded α-synuclein and constitute an efficient and previously underappreciated pathway for its intercellular transfer^32^. While these findings link sphingolipid dysregulation to neuron-to-neuron transmission of α-synuclein, they raise important questions regarding the role of astrocytes in this process. While astrocytes actively contribute to α-synuclein handling through its uptake and degradation, the extent to which astrocytes internalize neuron-derived ectosomes, and how this is modulated by disease-associated lipid alterations remains poorly understood.

Here, we sought to determine how astrocytes, in homeostatic versus disease-relevant states, differ in their capacity to internalize and process neuron-derived ectosomes. We find that GCase-deficient primary and PD patient iPSC-derived astrocytes exhibit impaired uptake of ectosome containing pathological α-synuclein. Mechanistically, this defect arises from compromised macropinocytosis. Importantly, the deficit in ectosome uptake driven by reduced GCase activity can be restored by enhancing lysosomal acidification or by limiting glucosylceramide synthesis. Together, these findings identify sphingolipid metabolism as a key regulator of ectosome internalization by astrocytes. Disruption of this pathway may impair astrocyte-mediated clearance of neuron-derived vesicles, thereby facilitating the extracellular persistence and propagation of α-synuclein pathology to neurons in PD.

## RESULTS

### GCase deficiency in astrocytes disrupts neuron-to-astrocyte lipid transport

Neuron-to-astrocyte intercellular transport is essential for maintaining brain homeostasis. Astrocytes are uniquely equipped to take up, store and/or degrade neuronal-derived cargos. Here we sought to test whether neuron-to-astrocyte transport is altered by GCase deficiency as is observed in PD. We first examined the transport of lipids as we have previously shown this pathway is disrupted in models of Alzheimer’s disease^33^.

Neurons were pulsed with BODIPY 558/568 C12 (Red-C12), a saturated 12-carbon fatty acid with a BODIPY 558/568 fluorophore covalently attached at the hydrophobic terminus, yielding a functional length comparable to an 18-carbon fatty acid. Red-C12 is widely used to study lipid trafficking and is incorporated into diverse lipid classes, including phospholipids and neutral lipids^34–36^. To assess loss of GCase activity, we either treated primary cortical cells with the irreversible GCase inhibitor conduritol-β-epoxide (CBE) (**Figure S1A-D**) or used dopaminergic neurons generated from human iPSCs carrying the N370S/+ mutation or their isogenic corrected control (**Figure S1E**)^37^. Both cortical and dopaminergic neurons internalize and store Red-C12 following an hour pulse (**Figure S1F-H**). While GCase-deficient neurons accumulate more Red-C12 puncta (**Figure S1F-H**), they also release more Red-C12 into the conditioned media compared to their respective controls (**Figure 1A-B**).

**Figure 1.**
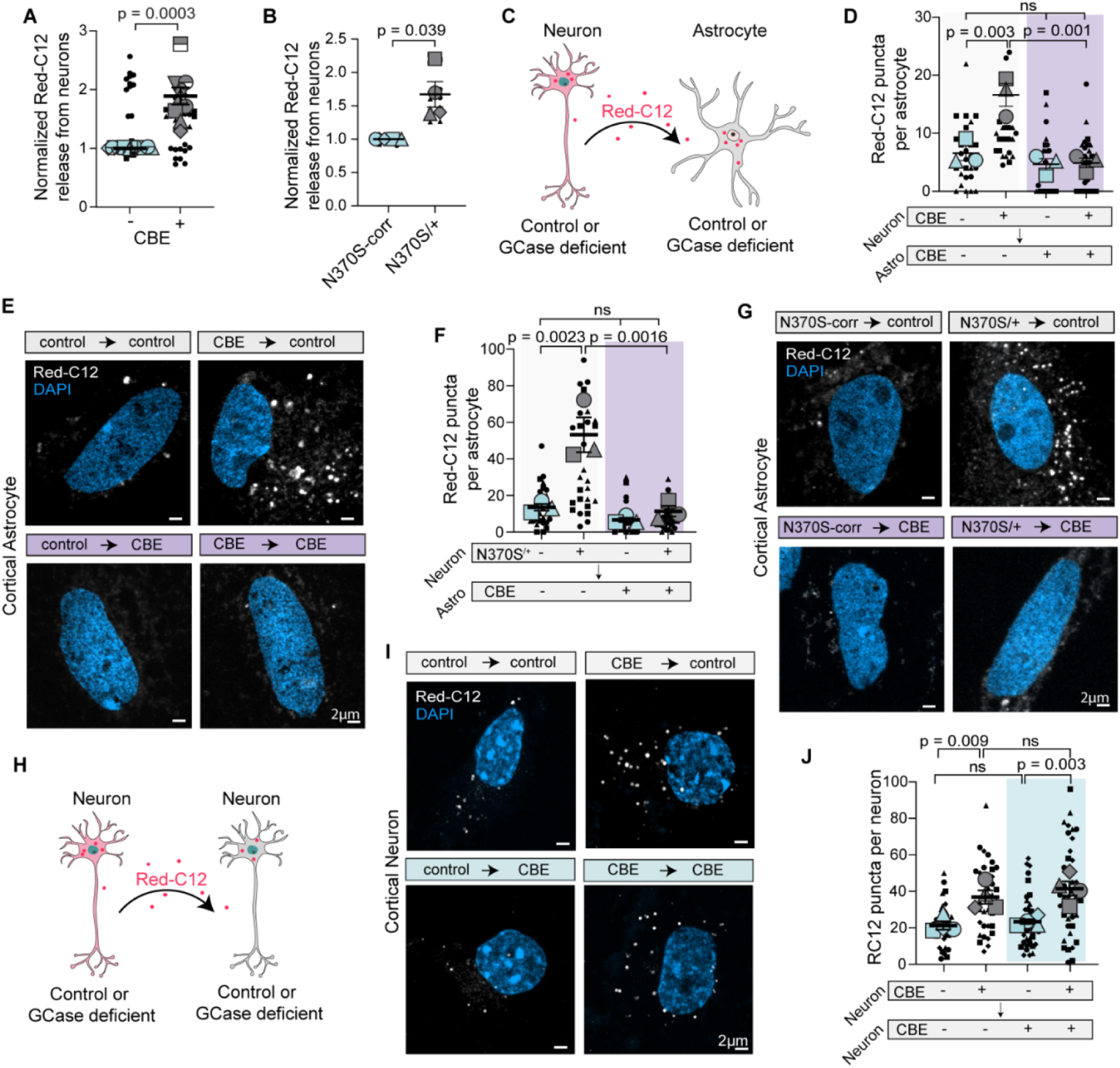
GCase deficiency impairs Red-C12 uptake in astrocytes but not neurons. A-B. Neuron-conditioned HBSS analyzed for Red-C12 fluorescence normalized to control neurons. n=4 independent experiments; 3 technical replicates/experiment; mean ± SEM. One sample *t*-test. C. Schematic depicting Red-C12 transfer assay from neurons to astrocytes. D-E. Red-C12 transfer from control or CBE-treated neurons to control or CBE-treated astrocytes. Maximum z-projections of astrocyte post-transfer assay. n = 4 independent experiments; 10 cells per experiment; mean ± SEM. One-way Anova with Šídák’s multiple comparisons test. F-G. Red-C12 transfer from control or GBA1-N370S/+ neurons to control or CBE-treated astrocytes. Maximum z-projections of astrocyte post-transfer assay. n = 4 independent experiments; 10 cells per experiment; mean ± SEM. One-way Anova with Šídák’s multiple comparisons test. H. Schematic depicting Red-C12 transfer assay from neurons to neurons. I-J. Red-C12 transfer from control or CBE-treated neurons to control or CBE-treated neurons. Maximum z-projections of astrocyte post-transfer assay. n = 4 independent experiments; 10 cells per experiment; mean ± SEM. One-way Anova with Šídák’s multiple comparisons test. Graphs are depicted as superplots where biological replicates are shown in large shapes, and technical replicates are shown as small shapes.

We next assessed the transfer of neuron-derived lipids to astrocytes. To this end, control or GCase deficient neurons pre-pulsed with Red-C12 overnight were incubated with control or GCase-deficient astrocytes on different coverslips separated by paraffin wax^38^. The appearance of Red-C12-positive puncta within astrocytes was quantified (**Figure 1C**). Consistent with increased lipid release from GCase-deficient neurons, control astrocytes exposed to CBE-treated cortical neurons or GBA1-N370S/+ dopaminergic neurons exhibited a marked increase in Red-C12–positive puncta compared to those exposed to control neurons (**Figure 1D-G**). Interestingly, this increase was abolished when recipient astrocytes were GCase-deficient (**Figure 1D-G)**. No difference in Red-C12 puncta size was observed (**Figure S1I-J**). In contrast, in neuron-to-neuron transfer assays (**Figure 1H**), GCase-deficient recipient neurons retained the ability to take up and accumulate Red-C12 puncta (**Figure 1I and 1J**), with no difference in Red-C12 puncta size (**Figure S1K**). We confirmed that neuron-derived Red-C12 accumulates in lipid droplets in astrocytes by showing 60% and 90% of the RedC12 puncta colocalize with Plin2 and BODIPY-493/503 respectively (**Figure S1L-O)**. Since storage of Red-C12 in lipid droplets is a requisite to detect Red-C12 puncta in these assays, we also wondered whether these results could be due to defects in lipid droplet formation. On the contrary, GCase deficiency increased the number of lipid droplets labelled with BODIPY-493/503 (**Figure S1P-Q**). Collectively, this data indicate a selective impairment in the uptake of neuron-derived lipids by GCase deficient astrocytes.

### GCase deficiency in astrocytes disrupts neuron-to-astrocyte α-synuclein transport

We next wondered whether GCase deficiency disrupts neuron-to-astrocyte transport of additional cargos. One physiologically relevant cargo in PD is α-synuclein. Intercellular transmission of misfolded α-synuclein has been proposed to play a key role in PD pathology^6,7,39^. We recently discovered α-synuclein is released within ectosomes and propagates α-synuclein pathology between PD neurons ^32^, and this neuron-to-neuron transport is increased in neurons with GCase deficiency. Since, approximately 70% of the ectosomes released by GCase-deficient dopaminergic neurons contain Red-C12 puncta (**Figure S2A-B**), and GCase deficiency impairs Red-C12 uptake, we wondered whether GCase deficiency also affects transport of ectosome from neurons to astrocytes.

To assess this, neurons expressing the membrane reporter mVenus-CAAX were preloaded with fluorescently tagged α-synuclein pre-formed fibrils (PFFs). In contrast to soluble monomers, these short fibrils, which are largely 50 nm in length (**Figure S2C-D**), are pathogenic due to their ability to induce the misfolding and aggregation of endogenous α-synuclein^40–42^. We assessed the transfer of mVenus-CAAX and α-synuclein PFF–Atto594 from control or GCase deficient neurons to control or GCase deficient astrocytes (**Figure 2A**). Like the Red-C12 transfer assays above, GCase deficiency, by either treating cortical neurons with CBE or using GBA1-N370S/+ dopaminergic neurons increased transfer of mVenus-CAAX and PFF-Atto594 to control astrocytes (**Figure 2B-G**). Strikingly, this increase was abolished when the recipient astrocytes were also GCase-deficient (**Figure 2B-G**). The size of mVenus-CAAX and α-synuclein PFF–Atto594 puncta were largely unaffected (**Figure S2E-H**). This data supports impaired neuron-to-astrocyte transport of α-synuclein with GCase deficiency.

**Figure 2.**
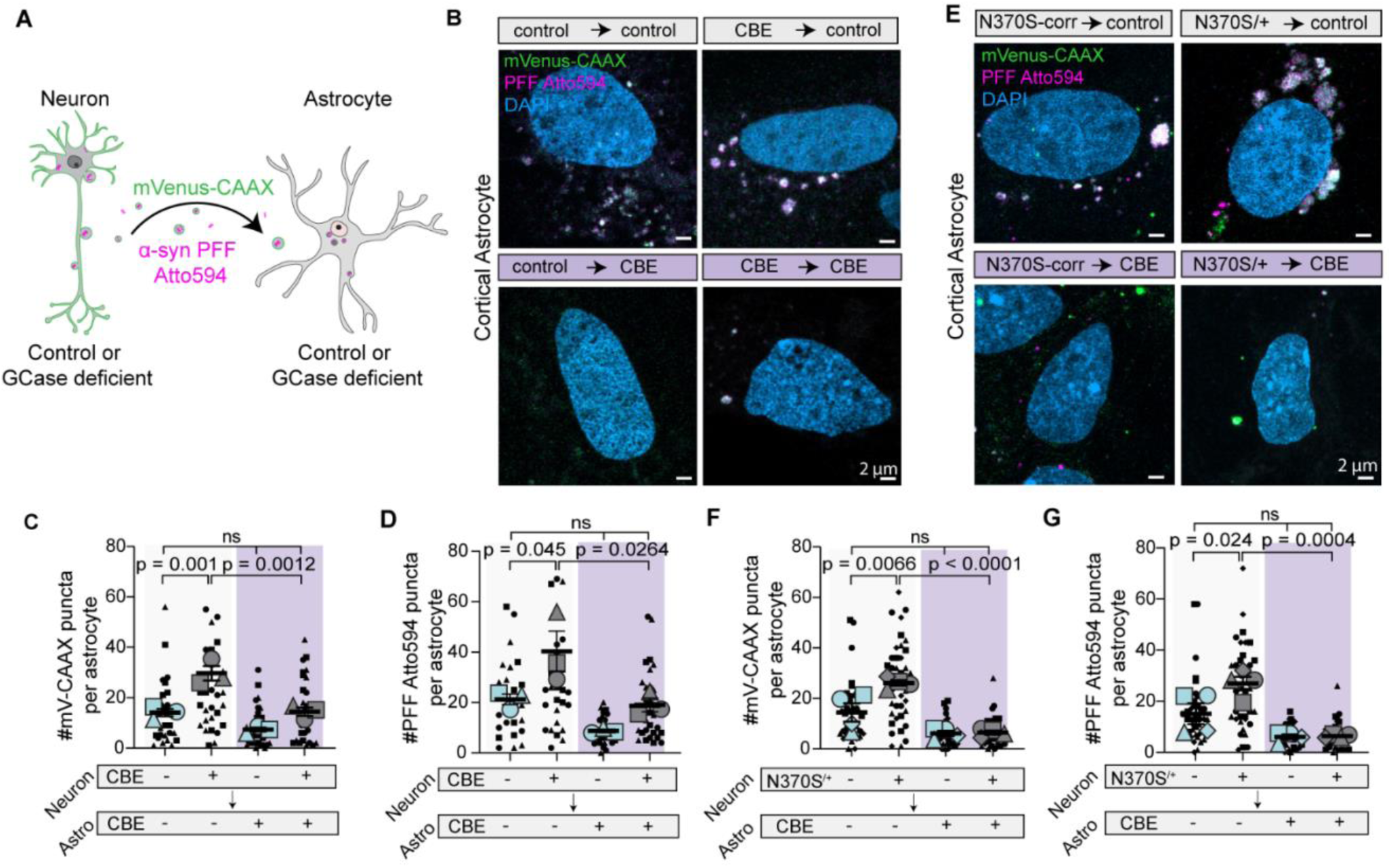
GCase deficiency impairs α-synuclein uptake by astrocytes. A. Schematic depicting transfer of mVenus-CAAX and PFF-ATTO-594 from neurons to astrocytes. B-D. mVenus-CAAX and PFF-ATTO594 transfer from control or CBE-treated neurons to control or CBE-treated astrocytes. n = 4 independent experiments; 10 cells per experiment; mean ± SEM. One-way Anova with Šídák’s multiple comparisons test. E-G. mVenus-CAAX and PFF-ATTO594 transfer from control or GBA1-N370S/+ neurons to control or CBE-treated astrocytes. n = 4 independent experiments; 10 cells per experiment; mean ± SEM. One-way Anova with Šídák’s multiple comparisons test. Graphs are depicted as superplots where biological replicates are shown in large shapes, and technical replicates are shown as small shapes.

### GCase deficiency reduces astrocytic uptake of ectosomes containing α-synuclein

With GCase deficiency, neurons preferentially release α-synuclein PFFs within ectosomes that are readily taken up by other neurons^32^. We therefore wondered whether GCase deficiency reduces the ability of astrocytes to take up ectosomes. To test this, ectosome-enriched fractions were prepared from control or GBA1-N370S/+ dopaminergic neurons expressing mVenus-CAAX and preloaded with α-synuclein PFF–Atto594 using differential centrifugation as previously established^32^. The presence of vesicles, roughly 1-2 μm in diameter, containing mVenus-CAAX and PFF-Atto594 in the fractions were confirmed (**Figure 3A**). These ectosome-enriched fractions were then applied to control or GCase-deficient cortical astrocytes (**Figure 3B**). As expected, GCase deficiency increased mVenus-CAAX and PFF-Atto594 puncta in control cortical astrocytes, whereas the uptake was greatly reduced in GCase-deficient astrocytes (**Figure 3C-E**). In addition, when treated with ectosomes from GBA1-N370S/+ neurons, puncta size of mVenus-CAAX and PFF-Atto594 were increased in control astrocytes, an effect that was absent in GCase-deficient astrocytes (**Figure S2I-J**).

**Figure 3.**
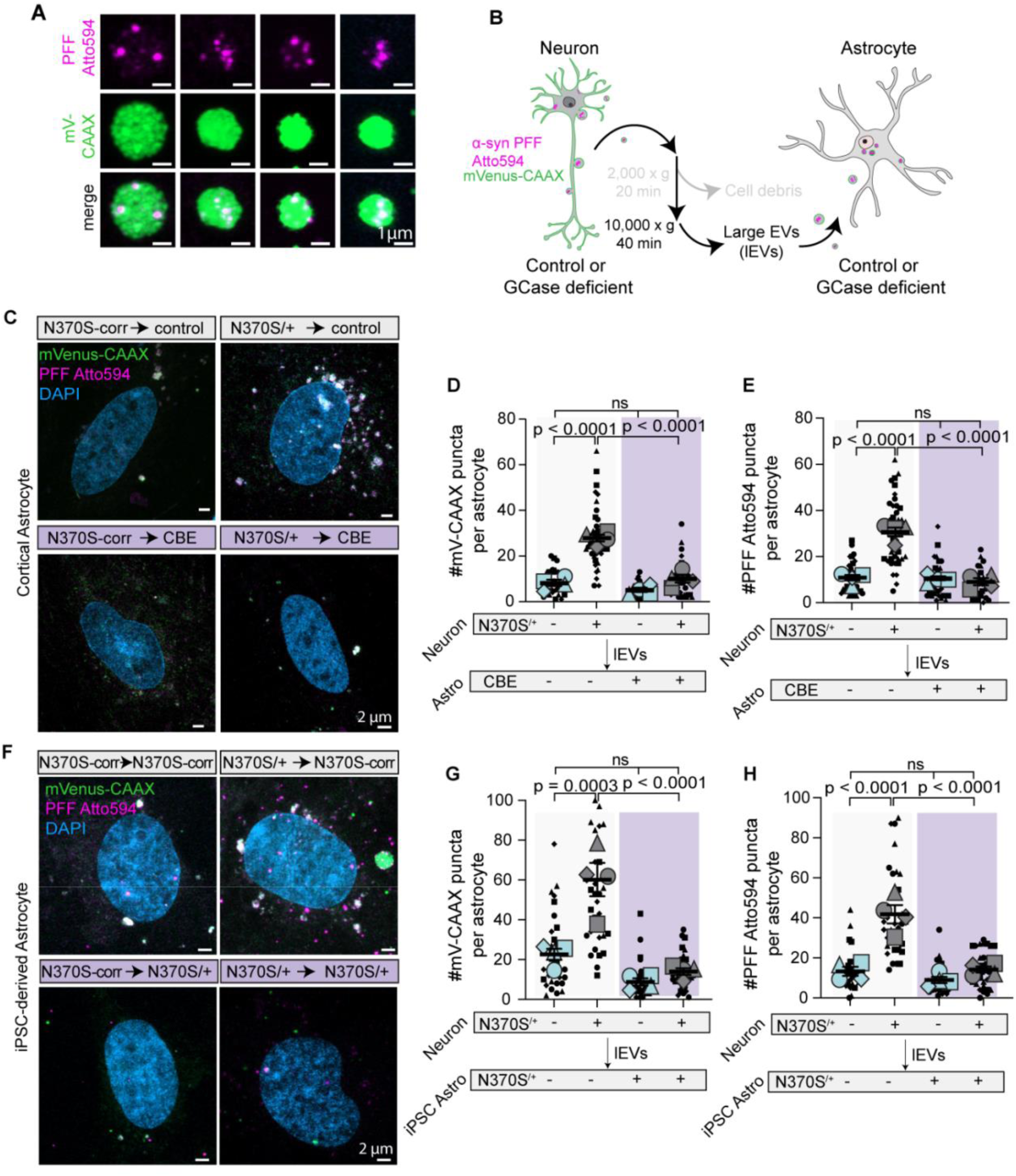
GCase deficiency affects ectosomal α-synuclein uptake by astrocytes. A. Maximum z-projections of mVenus-CAAX positive ectosomes containing PFF-ATTO-594 collected after differential centrifugation from neuron-conditioned media. B. Schematic depicting assay treating astrocytes with mVenus-CAAX and PFF-ATTO-594 containing ectosome fractions collected from neuron conditioned media. C-E. Control or CBE-treated astrocytes treated with mVenus-CAAX and PFF-ATTO594 containing ectosome fractions released by control or GBA1-N370S/+ neurons. n = 4 independent experiments; 10 cells per experiment; mean ± SEM. One-way Anova with Šídák’s multiple comparisons test. F-H. Control or GBA1-N370S/+ astrocytes treated with mVenus-CAAX and PFF-ATTO594 containing ectosome fractions released by control or GBA1-N370S/+ neurons. n = 4 independent experiments; 10 cells per experiment; mean ± SEM. One-way Anova with Šídák’s multiple comparisons test. Graphs are depicted as superplots where biological replicates are shown in large shapes, and technical replicates are shown as small shapes.

To determine whether this phenotype extends to human cells, we generated astrocytes from iPSCs derived from a PD patient carrying the N370S/+ mutation in *GBA1* and its corresponding isogenic control (**Figure S2K**). As expected, GBA1-N370S/+ astrocytes exhibited reduced GCase activity relative to its isogenic control (**Figure S2L**). As above isogenic control astrocytes accumulated more mVenus-CAAX and PFF-Atto594 puncta when treated with ectosome enriched fractions from GBA1-N370S/+ neurons (**Figure 3F-H**). However, this uptake was greatly reduced in GBA1-N370S/+ astrocytes (**Figure 3F-H**). The size of mVenus-CAAX and PFF-Atto594 puncta was unaffected (**Figure S2M-N**). Together, these experiments confirm that neuronal GCase deficiency enhances the release of lipids and α-synuclein–containing vesicles ^32^. More importantly, these experiments reveal that reduced GCase activity, as occurs in PD, prevents astrocytes from internalizing neuronal derived α-synuclein in ectosomes.

### Blocking glucosylceramide synthesis restores ectosomal α-synuclein uptake by astrocytes

GCase deficiency prevents the conversion of hexosylceramides (galactosylceramide and glucosylceramide) to ceramide, resulting in an accumulation of hexosylceramides^16,18,22,32^. These metabolic consequences of GCase inhibition were confirmed using untargeted lipidomics on control astrocytes and those treated with CBE. We identified 87 lipid species that increased and 53 that decreased with GCase deficiency (**Figure S3A**). All major lipid classes were affected, with sphingolipids representing the most prominently altered category (**Figure S3B**). Within this class, the most notable alteration was an accumulation of hexosylceramides (**Figure 4A**).

**Figure 4.**
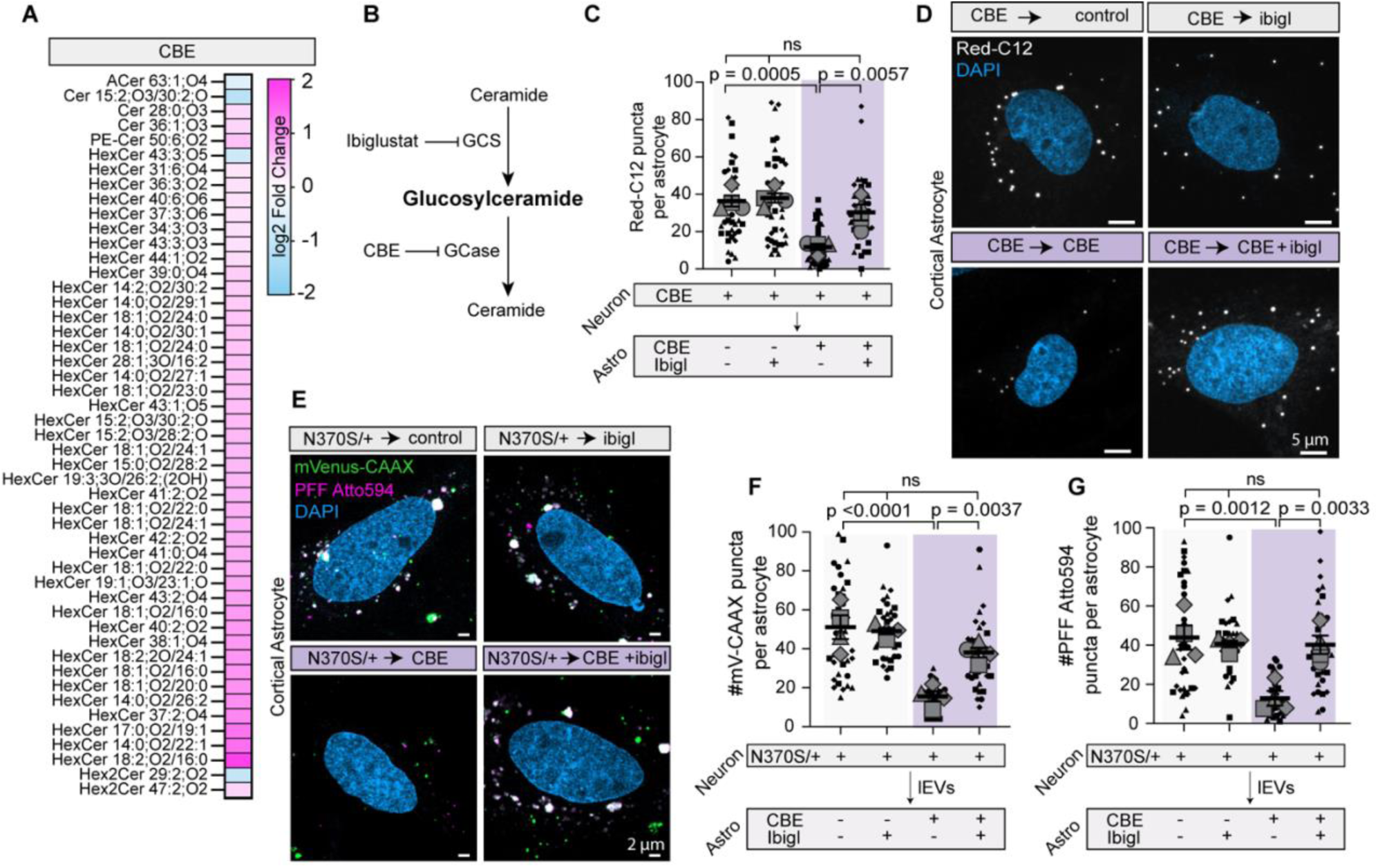
Blocking glucosylceramide synthesis recovers Red-C12 and ectosomal α-synuclein uptake by astrocytes. A. Heatmap of altered sphingolipid species (fold change (FC) > 1.40 or < 0.71 and p < 0.05) upon CBE treatment in cortical astrocytes. n = 4 independent experiments; ceramides (Cer), galactosyl- and glucosylceramides (HexCer) and lactosylceramides (Hex2Cer). B. Schematic of key enzymes in glucosylceramide synthesis and hydrolysis. Glucosylceramide synthase (GCS). C-D. Red-C12 transfer from CBE neurons to control, CBE, and/or ibiglustat-treated astrocytes. Maximum z-projections of astrocyte post-transfer assay. n = 4 independent experiments; 10 cells per experiment; mean ± SEM. One-way Anova with Šídák’s multiple comparisons test. E-G. Control, CBE and/or ibiglustat-treated astrocytes treated with mVenus-CAAX and PFF-ATTO594 containing ectosome fractions released by GBA1-N370S/+ neurons. Maximum z-projections of astrocytes post-treatment. n = 4 independent experiments; 10 cells per experiment; mean ± SEM. One-way Anova with Šídák’s multiple comparisons test. Graphs are depicted as superplots where biological replicates are shown in large shapes, and technical replicates are shown as small shapes.

We previously found that increased glucosylceramide is responsible for ectosome shedding from neurons^32^. Since our lipidomics analysis could not distinguish hexosylceramides as galactosylceramide or glucosylceramide, we sought to determine whether impaired astrocytic cargo uptake was also due to increased glucosylceramides. To test this, we inhibited glucosylceramide synthase using Ibiglustat (**Figure 4B**)^22,43^. First, Red-C12 transfer assays were performed (**Figure 1C**). As above, GCase deficiency in astrocytes induced by CBE treatment decreased the accumulation of neuron-derived Red-C12 (**Figure 4C-D**). Preventing glucosylceramide synthesis with ibiglustat restored the number of Red-C12 puncta, with no change in puncta size (**Figure 4C-D and S3C**). We next tested the uptake of neuron-derived ectosomes containing mVenus-CAAX and PFF-Atto594 as above (**Figure 3B**). Again, while GCase deficiency reduced uptake of ectosomes containing α-synuclein, this was rescued by co-treating astrocytes with ibiglustat (**Figure 4E-G**). Here, we again noted a reduction in the size of mVenus-CAAX and PFF-Atto594 puncta in CBE-treated astrocytes which was restored with ibiglustat (**Figure S3D-E**). In these assays, while ibiglustat alone did not affect cytotoxicity, cytotoxicity was increased when combined with CBE and ectosomes containing α-synuclein (**Figure S3F**). These data indicate that increased glucosylceramide is responsible for reducing the ability of astrocytes to take up neuron-derived ectosomes containing α-synuclein.

### Improving lysosomal function in astrocytes enhances Red-C12 and ectosomal α-synuclein uptake

As GCase is a lysosomal enzyme^15^, and both CBE and mutations in *GBA1* cause lysosomal dysfunction^18,44^, we wondered whether restoring lysosomal function could rescue astrocytic capacity for uptake. To this end, GCase-deficient astrocytes were treated with rimeporide to inhibit the sodium/hydrogen exchanger type 1^45^. This approach has previously been used to reacidify lysosomes and recover lysosomal function in *GBA1* and *LRRK2* mutant dopaminergic neurons and in cells lacking progranulin, another PD-related gene^32,46^. We first assessed LysoTracker as a readout of functional acidic compartments. Consistent with previous work, LysoTracker staining and number of puncta were reduced following CBE treatment and restored to levels comparable to controls when co-treated with rimeporide (**Figure 5A-C**). We then tested if restoring lysosomal acidity could rescue disrupted uptake by GCase-deficient astrocytes. Indeed, rimeporide rescued both the number of Red-C12 puncta in the lipid transport assay (**Figure 5D-E**), and the number and size of mVenus-CAAX and PFF-Atto594 puncta when treated with neuron-derived ectosomes (**Figure 5F-H and S3H-I**), without inducing cytotoxicity (**Figure S3J**). Together, these data demonstrate that restoring lysosomal acidification may represent a promising strategy to recover the ability of astrocytes to internalize neuron-derived cargos.

**Figure 5.**
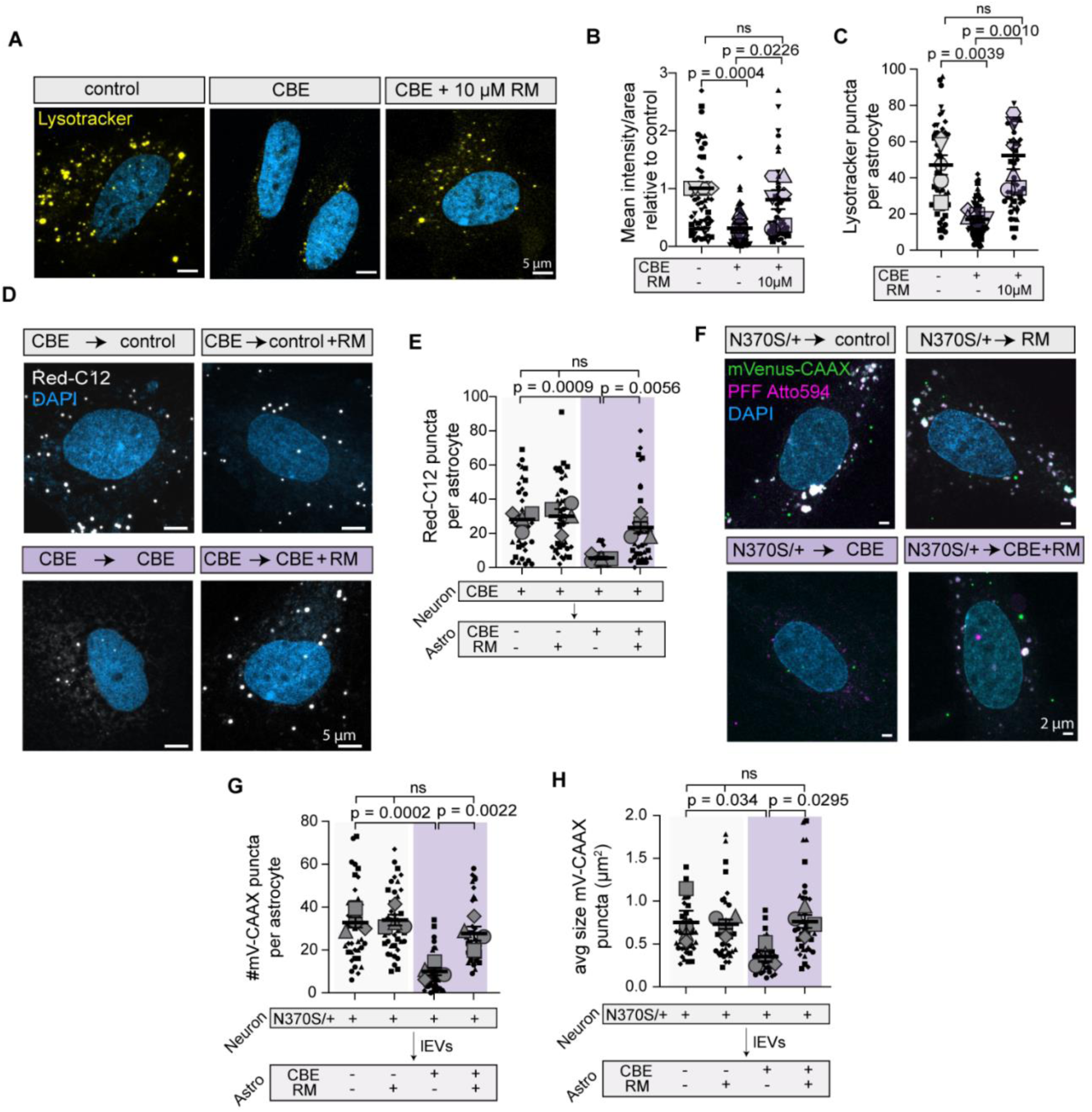
Enhancing lysosomal function recovers uptake of lipids and ectosomal α-synuclein by GCase-deficient astrocytes. A-C. Maximum intensity projections of control and GCase-deficient astrocytes +/- rimeporide (RM), stained with lysotracker. Quantification of lysotracker intensity and number of puncta. n = 4 independent experiments; 10 cells per experiment; Mean ± SEM; One-way ANOVA with Šidák’s correction for multiple comparisons. D-E. Red-C12 transfer from CBE-treated neurons to control, CBE, and/or rimeporide-treated astrocytes. Maximum z-projections of astrocyte post-transfer assay. n = 4 independent experiments; 10 cells per experiment; mean ± SEM. One-way Anova with Šídák’s multiple comparisons test. F-H. Control, CBE and/or rimeporide-treated astrocytes treated with mVenus-CAAX and PFF-ATTO594 containing ectosome fractions released by GBA1-N370S/+ neurons. Maximum z-projections of astrocytes post-treatment. n = 4 independent experiments; 10 cells per experiment; mean ± SEM. One-way Anova with Šídák’s multiple comparisons test. Graphs are depicted as superplots where biological replicates are shown in large shapes, and technical replicates are shown as small shapes.

### Reduced macropinocytosis accounts for reduced ectosomal α-synuclein uptake in GCase deficient astrocytes

Astrocytes could internalize extracellular vesicles through multiple pathways such as endocytosis, phagocytosis, and macropinocytosis. Given the reduction in astrocytic uptake of ectosomes with GCase deficiency, we next sought to identify which of these pathways is most affected by impaired GCase activity. To this end, we pharmacologically inhibited distinct uptake routes in astrocytes using Pitstop2, an inhibitor of endocytosis, or EIPA, which disrupts macropinocytosis^47–49^. We confirmed the specificity of these drugs by showing that Pitstop2, but not EIPA, blocked transferrin uptake, an endocytic cargo (**Figure S4A-B**). Conversely EIPA, but not Pitstop2, blocked the uptake of a high molecular weight dextran known to be taken up by macropinocytosis (**Figure S4C-F**). In these assays, while EIPA did not affect cytotoxicity, cytotoxicity was increased using Pitstop2 (**Figure S4G**).

We then assessed if the internalization of neuron-derived cargo is affected by these inhibitors. First, EIPA but not Pitstop2, reduced the amount of neuron-derived Red-C12 in astrocytes without affecting puncta size (**Figure 6A-D and S4H-I**). While Red-C12 continues to be internalized in the presence of Pitstop2, this treatment also alters the morphology of Red-C12 into what appears to be enlarged endosomal structures similar to those previously reported (**Figure 6C**)^50^. We next tested whether blocking macropinocytosis similarly blocked uptake of ectosomes. Indeed, EIPA, reduced the number and size of mVenus-CAAX and PFF-Atto594 in astrocytes treated with neuron-derived ectosomes (**Figure 6E-G and S4J-K**). Pitstop2, on the other hand, had no effect on the size or number of mVenus-CAAX and PFF-Atto594 (**Figure S4L-P).**

**Figure 6.**
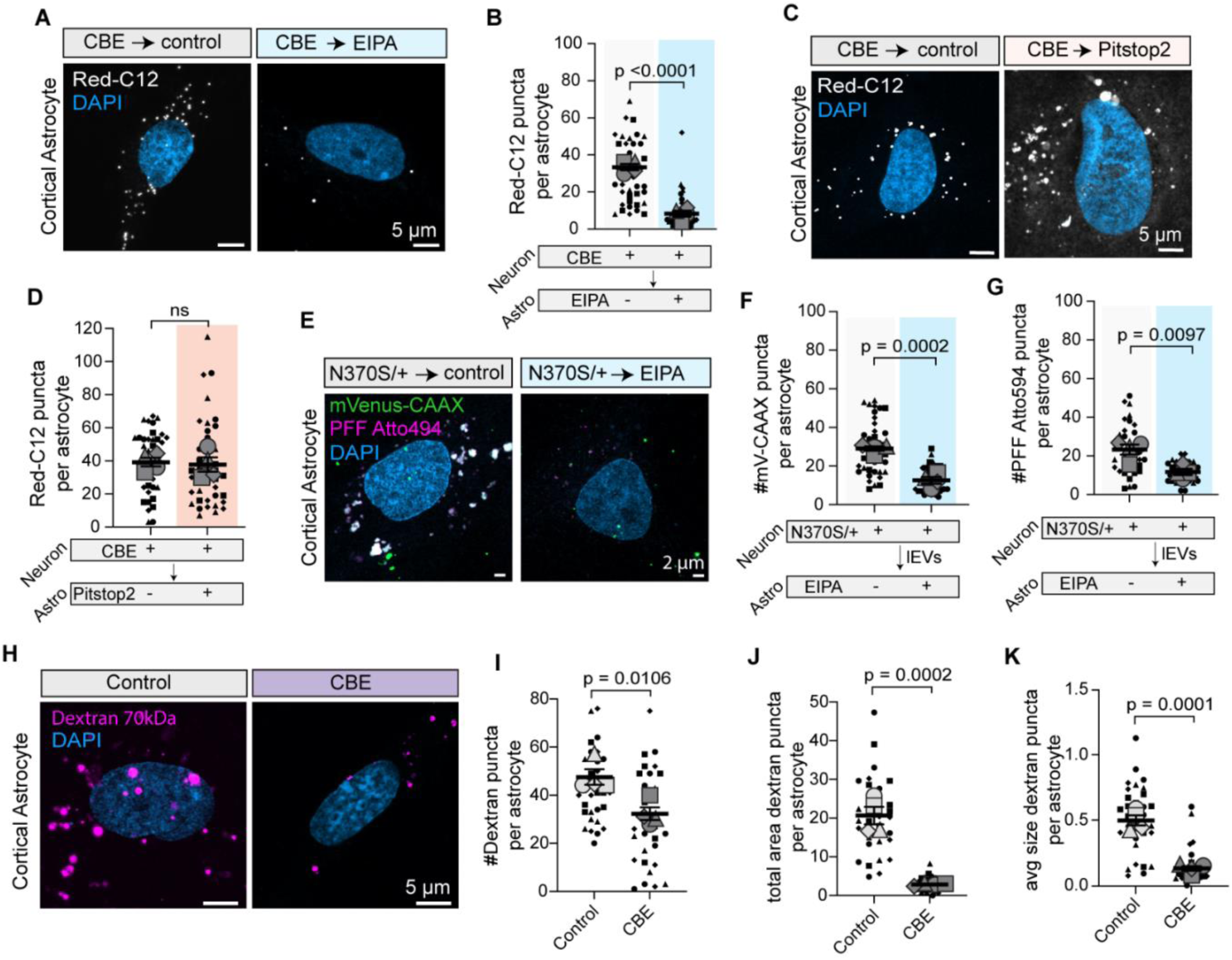
Impaired macropinocytosis in GCase-deficient astrocytes reduces uptake of neuronal lipids and ectosomes containing α-synuclein. A-B. Red-C12 transfer from CBE-treated neurons to control or EIPA-treated astrocytes. Maximum z-projections of astrocyte post-transfer assay. n = 4 independent experiments; 10 cells per experiment; mean ± SEM. One-way Anova with Šídák’s multiple comparisons test. C-D. Red-C12 transfer from CBE-treated neurons to control or Pitstop2-treated astrocytes. Maximum z-projections of astrocyte post-transfer assay. n = 4 independent experiments; 10 cells per experiment; mean ± SEM. One-way Anova with Šídák’s multiple comparisons test. E-G. Control, or EIPA-treated astrocytes treated with mVenus-CAAX and PFF-ATTO594 containing ectosome fractions released by GBA1-N370S/+ neurons. Maximum z-projections of astrocytes post-treatment. n = 4 independent experiments; 10 cells per experiment; mean ± SEM. One-way Anova with Šídák’s multiple comparisons test. H-L. Control, or CBE-treated astrocytes treated with Texas-Red labelled high molecular weight dextran (70 kDa). Maximum z-projections of astrocytes post-treatment. n = 4 independent experiments; 10 cells per experiment; mean ± SEM. Unpaired *t*-test. Graphs are depicted as superplots where biological replicates are shown in large shapes, and technical replicates are shown as small shapes.

Finally, we reasoned that if ectosomes are internalized by macropinocytosis, and GCase deficiency reduces ectosome internalization, that this is a result of reduced macropinocytosis. To test this, we treated control or GCase deficient astrocytes with different cargos as above. GCase deficient astrocytes had decreased uptake of the macropinocytosis cargo high molecular weight dextran (**Figure 6H-K**) and no effect on the endocytic cargo transferrin (**Figure S4Q-R**). Collectively, these data indicate that astrocytic uptake of neuron-derived ectosomes occurs via macropinocytosis and this uptake is disrupted by GCase deficiency.

## DISCUSSION

PD is a multicellular disorder involving complex interactions between neurons and glial cells^4,5^. Neuron-to-astrocyte intercellular transport is essential for maintaining brain homeostasis, as astrocytes are uniquely equipped to take up, store, and degrade neuronal-derived cargos^28,29,33,51^. Although astrocytes are increasingly recognized as important regulators of extracellular protein homeostasis^33,52^, most studies have focused on their capacity to internalize free α-synuclein species^47^. Our findings extend this concept by demonstrating that astrocytes, under normal conditions, efficiently internalize ectosomes containing α-synuclein released by neurons by macropinocytosis. However, when GCase activity was reduced, as occurs in PD astrocytes lose this capacity. This provides an important mechanism by which GCase deficiency could reduce the ability of astrocytes to clearance misfolded protein in disease.

Here, we identified glucosylceramide accumulation as a major contributor to impaired astrocytic uptake. GCase deficiency produced widespread lipid remodeling, with sphingolipids representing the most prominently altered lipid class. Pharmacological inhibition of glucosylceramide synthase restored both lipid transfer and uptake of α-synuclein-containing ectosomes, implicating excess glucosylceramide as a key mediator of the phenotype. These findings complement previous work demonstrating that glucosylceramide accumulation promotes α-synuclein aggregation and stimulates ectosome shedding from neurons^32^. Together, these studies suggest that glucosylceramide dysregulation contributes to multiple stages of α-synuclein propagation, simultaneously enhancing vesicle release from donor neurons while impairing clearance by recipient astrocytes. Such a dual mechanism would be expected to amplify the spread of pathological α-synuclein throughout neural circuits.

The ability of rimeporide, an NHE-1 inhibitor, to rescue ectosome uptake is consistent with the importance of lysosomes in regulating cargo internalization. Lysosomal dysfunction is a well-established consequence of GCase deficiency and a common feature of both familial and sporadic forms of PD^18,20,27,53^. Consistent with PD neurons^18,20,54^, lysosomal acidification in GCase-deficient astrocytes is restored by rimeporide treatment. In addition to regulating lysosomal acidity, NHE-1 is present on the plasma membrane and influences cellular acidity. Whether increased cellular acidity contributes to recovery of ectosome uptake in rimeporide-treated astrocytes remains unclear. Given that the non-specific NHE inhibitor amiloride can reduce macropinocytosis by increasing localized acidity on the plasma membrane ^47–49^, we anticipate the results here are likely due to the regulation of the lysosome.

An important finding of this study is the identification of macropinocytosis as the mechanism through which astrocytes internalize neuron-derived ectosomes. Our results also indicate that GCase deficiency selectively disrupts macropinocytosis rather than causing a generalized defect in endocytosis. Macropinocytosis has previously been implicated in the uptake of protein aggregates and extracellular vesicles by glial cells^47,55^, but much remains unknown regarding its regulation by sphingolipid metabolism. Our results suggest that altered membrane composition resulting from glucosylceramide accumulation may impair the membrane remodeling events required for efficient macropinosome formation.

The observation that neuron-derived ectosomes containing α-synuclein are preferentially taken up by astrocytes under physiological conditions raises important implications for disease progression. Astrocytes are generally considered more resistant than neurons to α-synuclein toxicity and possess robust degradative systems capable of processing extracellular protein aggregates^56^. Consequently, astrocytic uptake of ectosomes may normally represent a protective sink that limits neuronal exposure to pathogenic α-synuclein species. In the context of GCase deficiency, however, this protective mechanism is compromised. Combined with our previous finding that GCase-deficient neurons release greater numbers of α-synuclein-containing ectosomes^32^, the present data support a model in which disease-associated lipid dysregulation simultaneously increases ectosome production by neurons and decreases astrocytic uptake. Consistent with this scenario, we found that GCase deficiency decreased neuronal lipid transport from neurons to astrocytes, but neuron to neuron transport was unaffected. Therefore, the resulting accumulation of extracellular vesicles in PD could enhance neuron-to-neuron transmission of α-synuclein and accelerate disease spread. The selective nature of this defect is noteworthy, as they support the possibility that astrocytes and neurons employ distinct uptake mechanisms for extracellular vesicles and that astrocytic pathways are uniquely sensitive to lipid disruptions as occurs in PD^29,57^. Such cell-type specificity may help explain why astrocytes, which are central regulators of lipid metabolism in the brain^58^, are disproportionately affected by perturbations in *GBA1*-associated pathways. More broadly, these results emphasize that the consequences of GCase deficiency extend beyond neuronal dysfunction and involve critical alterations in neuron–glia communication.

In summary, this study demonstrates that GCase deficiency selectively impairs astrocytic uptake of neuron-derived ectosomes containing α-synuclein through disruption of macropinocytosis. This defect is driven by altered sphingolipid metabolism and lysosomal dysfunction and can be rescued by reducing glucosylceramide synthesis or restoring lysosomal acidification. These findings reveal a previously unrecognized role for astrocytes in regulating ectosome-mediated α-synuclein trafficking and suggest that loss of astrocytic clearance mechanisms may contribute to the propagation of pathology in PD. Targeting sphingolipid metabolism may therefore represent promising therapeutic strategies to restore uptake by astrocytes and limit the spread of α-synuclein pathology.

### Limitations of the study

In this study, we provide compelling evidence that GCase deficiency reduces the internalization of ectosomes containing α-synuclein by astrocytes. Nevertheless, some limitations of the study are worth noting. First, the study primarily relies on in vitro primary and iPSC-derived cell models, which may not fully recapitulate the complexity of the brain. Interactions with microglia, oligodendrocytes, and the extracellular matrix may influence vesicle trafficking and uptake in ways not captured here. In addition, while our findings identify glucosylceramide accumulation and lysosomal dysfunction as important contributors, the precise molecular events linking these alterations to impaired macropinocytosis remain to be elucidated. Finally, whether similar defects occur in sporadic PD or other forms of synucleinopathies characterized by reduced GCase activity will require further investigation.

## METHODS

### Animals

Experiments involving rat cells were approved by and performed under the Canadian Council of Animal Care at the University of Alberta (AUP#3358). Sprague-Dawley-timed pregnant rats were obtained from Charles River Laboratories and arrived at our facility 1 week before birth.

### Primary culture of cortical neurons

Mixed sex primary cortical cultures were prepared as previously described for hippocampal cultures^38^. Briefly, cortices were dissected from P0-P1 Sprague-Dawley rat pups and digested with papain (Fisher Scientific #LK003178) and Benzonase® Nuclease (Sigma #E1014). After digestion, the tissue was gently triturated and filtered with a 70-µm nylon cell strainer (Fisher Scientific #22363548). Neurons were grown on poly-D-lysine (Sigma-Aldrich, #P6407) coated glass coverslips or plastic tissue culture dishes in neurobasal medium (Fisher Scientific, #21103049) containing B-27 supplement (Fisher Scientific #17504044), 2 mM GlutaMax (Fisher Scientific #35050061) and antibiotic-antimycotic (Fisher Scientific #15240062). 2 μM AraC was added to the media 2 days after plating to prevent the contamination of glial cells in neuronal cultures. Primary cortical astrocytes were grown in Basal Eagle Media containing 10% fetal bovine serum, 0.45% glucose, 1 mM sodium pyruvate (Fisher Scientific #11360070), 2 mM GlutaMax (Fisher Scientific #35050061) and antibiotic-antimycotic (Fisher Scientific #15240062). All cells were grown at 37°C in 5% CO2 and used at DIV 7.

### iPSC-derived dopaminergic neurons and astrocytes

The use of human iPSCs in the EDDU was approved by the McGill University Research Ethics Board (IRB Study Number A03-M19-22A). Human iPSC lines derived from PD patients with mutations in GBA1 [N370S (female/65 yrs)^37^, and its isogenic control were obtained. Characterization and quality control data regarding the GBA1 iPSCs are provided in^32^ and on iPSCRg website. iPSCs were induced into dopaminergic neurons and astrocyte progenitors as described previously in^37,59,60^. CRISPR editing of patient-derived iPSCs to generate iPSCs was performed according to earlier published workflows^37,61^.

Dopaminergic neural progenitor cells were dissociated with StemPro Accutase Cell Dissociation Reagent (ThermoFisher #A1110501) into single-cell suspensions. Cells were plated onto laminin (Fisher Scientific #23017015) and poly-L-ornithine (Sigma #P3655) coated coverslips in 6-well plates (1×105 cells) for immunostaining and live-cell imaging or on coated 10 cm dishes (1×106 cells) for vesicle collection by differential ultracentrifugation, with neural progenitor plating medium (DMEM/F12 supplemented with N2, B27 supplement; Fisher Scientific #11320082, 17502048, 17504044). Progenitors were differentiated into dopaminergic neurons by switching to differentiation medium (Fisher Scientific #21103049) supplemented with N2 and B27, BDNF (20 ng/mL; Peprotech #450-02), GDNF (20 ng/mL; Peprotech #450-10), Compound E (0.1 μM; Stemcell Technologies #73954), db-cAMP (0.5 mM; Cedarlane # ND07996), Ascorbic acid (200 μM; Sigma #A5960), TGF-β3 (10 ng/mL; Peprotech #100-63E) and laminin (1 μg/mL; Thermofisher #23017015). Mitomycin C (0.1μg/mL; Fisher Scientific, #AAJ63193MA) was added for 7 days to prevent glial contamination.

Astrocyte progenitors, generated as described previously^60^ and positive for SOX2 and PAX6, were seeded at low cell density (15,000 cells/cm2) in two T25 flasks in the presence of 5ml of NPC expansion medium [DMEM/F12 medium supplemented with 1% N2, 2% B27, 20 ng/ml EGF (PeproTech; Cat. No. AF-100-15), 20 ng/ml FGF2 (PeproTech; Cat. No. 450-18b), 1 mM GlutaMax, 1% non-essential amino acids, 1 μg/ml laminin and antibiotic-antimycotic (Fisher Scientific #15240062)] containing Rock inhibitor Y-27632. The following day, medium was replaced with ‘Astrocyte Differentiation Medium 1’ [ScienCell Astrocyte Growth Medium (ScienCell Research Laboratories; Carlsbad, CA, USA; Cat. No. 1801b), astrocyte growth supplement (1x, starting from a 100x stock, as specified by the supplier, ScienCell Research Laboratories; Cat. No. 1852), 2% fetal bovine serum (FBS) (ScienCell Research Laboratories; Cat. No. 0010), 50 U/ml penicillin G, 50 mg/ml streptomycin]. Half medium was replaced with fresh medium every 3 to 4 days and cells were maintained under these culture conditions for 30 days without passaging. When plated for experiments astrocytes were maintained in Astrocyte Differentiation Medium 1 supplemented with 10% FBS.

### In vitro drug treatments

Conduritol-β-epoxide (CBE, Cayman #15216, MedChem Express #HY-100944) was reconstituted at 100 mM in dimethyl sulfoxide (DMSO) and stored at −20°C. Primary cortical neurons and astrocytes were treated overnight at DIV6 with 100μM final concentration prior to use. Ibiglustat (MCE no. HY-16743) was reconstituted at 10 mM in DMSO and stored at −80 °C. Astrocytes were treated O/N at DIV6 with a final concentration of 1μM. The specificity of ibiglustat has been independently verified ^22,43^. Rimeporide (Federline no. 332-12602-2) was reconstituted at 10 mM in DMSO and stored at −80 °C. To assess reacidification, astrocytes were treated with 10 μM rimeporide. A total of 100 nM Lysotracker Red DND-99 (Fisher no. L7528) was added to the neuron-conditioned media for 30 min at 37 °C, washed twice with warm PBS and fixed in 4% paraformaldehyde (PFA). EIPA (5-(N-Ethyl-N-isopropyl) amiloride) (Milipore Sigma, A3085) was reconstituted at 20mM in DMSO and stored at −80 °C. Astrocytes were treated at DIV7 with a final concentration of 20μM for 30 minutes in HBSS. Pitstop2 (Sigma, SML 1169), was reconstituted at 25mM in DMSO and stored at −80 °C. Astrocytes were treated at DIV7 with a final concentration of 25μM for 30 minutes in HBSS. Alexa Fluor® 647 - conjugated ChromPure Rat Transferrin (Cedarlane 012-600-050) was reconstituted in 2 mg/mL DMSO and stored at −20 °C. Astrocytes were treated at DIV7 with a final concentration of 0.5 μg/mL for 1 hour in HBSS. Dextran Texas Red 70 kDa (Thermo Fisher Scientific, D1864) was reconstituted in 10 mg/mL DMSO and stored at −20 °C. Astrocytes were treated at DIV7 with a final concentration of 0.2 mg/mL for 1 hour in HBSS.

### α-synuclein fibril preparation and characterization

ATTO 594 α-synuclein PFFs (Stressmarq #SPR-322E-A594) were generated according to The Michael J. FOX Foundation protocol. Prior to use, PFFs were diluted in PBS and sonicated (30s, 0.5s on/off at 30% power) using a UP50H ultrasonic processor. Negative stain electron microscopy: 100 μg/mL PFFs before and after sonication were added to glow discharged 200 copper mesh carbon grid, washed with ddH_2_O and stained with 3% uranyl acetate (Electron Microscopy Sciences #22400) for 1 min. PFFs were visualized using a 200 kv JEOL 2100 Transmission Electron Microscope and seed length was determined with Fiji-ImageJ2 and the skeleton plugin. To treat neurons, PFFs diluted to 2 μg/mL were added to cortical neurons (DIV7) or dopaminergic neurons 7 days post final differentiation, incubated for 24 h and used at 7 days post-treatment as indicated.

### Lentivirus production and transduction

Lentiviral plasmids for mVenus–CAAX was generated as previously described ^32^. Viral particles were produced in HEK293T cells as previously described. In brief, HEK293T cells were transfected with lentiviral plasmids by calcium phosphate transfection. Media containing viral particles 24 h, 36 h and 48 h post transfection were collected and filtered with a 0.45-μm filter to remove cell debris and concentrated by centrifugation at 7,441g for 16 hours at 4 °C. Cortical and dopaminergic neurons were transduced with lentivirus (MOI 2) on DIV3. The media were replaced with half-fresh half-conditioned cortical and dopaminergic cell media after 3 h, and the cells were used on DIV7.

### Collection of large extracellular vesicles, ectosomes

The neuron-conditioned medium of dopaminergic neurons was collected on day 14 of the final differentiation. The media was then centrifuged at 2,000 × g at 4 °C for 20 min, the resulting supernatant was centrifuged at 10,000 × g at 4 °C for 40 min. For treatment use, pellets were resuspended in ice-cold PBS and given to dopaminergic neurons, cortical astrocytes or human-iPSC derived astrocytes on the day of their final differentiation and incubated for 4 hours in HBSS.

### Lipidomic analyses

At DIV6 cortical astrocytes were treated with DMSO or 100 µM CBE. On DIV7, cells were washed twice in ice-cold PBS, scraped into ice-cold 1:1 methanol:water, and stored in liquid nitrogen. Lipidomic analysis from 5 independent experiments per group by the Metabolomic Innovation Center. In brief, samples were spiked with NovaMT LipidRep Internal Standard Basic Mic for Tissue/Cells containing 15 deuterated lipids, dichloromethane, and methanol. Lipids were extracted using a modified Folch liquid-liquid extraction protocol, air dried, resuspended in NovaMT MixB and diluting in NovaMT MixA, LC-MS analyses were performed on the lipid extracts using Thermo Vanquish UHPLC linked to Bruker Impact II QTOF Mass Spectrometer. Data of samples were independently acquired in both positive and negative ionization modes with mass range m/z 150-1500. Data post-processing and normalization were performed using LipidScreener (Nova Medical testing Inc.).

### Red-C12 and mVenus-CAAX ATTO594-PFF transfer assays

Primary cortical neurons and iPSC-derived dopaminergic neurons were incubated in complete medium supplemented with 4 μM Red-C12 for 16 hours, followed by two washes with warm HBSS and subsequently treated in HBSS for 4 hours at 37 °C. Red-C12–labeled neurons, or neurons expressing mVenus-CAAX and treated with ATTO594-PFF (for 7days), as well as unlabeled glial cells or neurons, were grown on separate coverslips. After washing twice with warm HBSS, the coverslips were placed face-to-face, separated by a thin layer of paraffin wax, and incubated in HBSS for 4 hours at 37 °C. Subsequently, the recipient cells were fixed with 4% warm PFA. Imaging was performed using a Zeiss LSM900 confocal microscope. For quantification, three-dimensional image stacks were converted to maximum intensity projections and processed using the RenyiEntropy thresholding algorithm. The number of Red-C12 or mVenus-CAAX/ATTO594-PFF–positive puncta were quantified in Fiji (ImageJ). Puncta localized within cells were identified by adjusting image brightness and contrast to clearly delineate cell boundaries. Previous analyses have shown that Red-C12 leakage from coverslips or wax is minimal and does not significantly affect the assay.

### Red-C12 release assays

Primary cortical neurons or iPSC-derived dopaminergic neurons were cultured on poly-D-lysine (PDL)- coated six-well plates and incubated with or without 4 μM Red-C12 for 16 hours in complete medium. Following incubation, cells were washed three times with warm PBS, incubated in fresh medium for 1 hour, and subsequently treated in HBSS for 4 hours at 37 °C. After treatment, the neuron-conditioned medium was collected and centrifuged at 2,000 × g for 10 minutes to eliminate dead cells. Supernatants were analyzed for Red-C12 using a Synergy Mx Multi-Mode Microplate Reader (BioTek Instruments Inc.). For each experiment technical triplicates were performed, and the results were averaged for quantification.

### Red-C12 accumulation assays

Cortical and iPSC-derived dopaminergic neurons were exposed to complete medium containing 4 μM Red-C12 for 30 minutes at 37 °C, followed by three washes with warm PBS and incubation in fresh neuronal medium for an additional 30 minutes at 37 °C. Cells were then washed twice with warm PBS. After treatment, neurons were fixed with warm 4% PFA and imaged using a Zeiss LSM900 Quantification was performed as described for the transfer assay.

### MTT assay

Cells (50,000 cells per well) were plated in 96-well flat-bottom plates and subjected to 1μM Ibiglustat (MCE no. HY-16743), α-synuclein PFFs-Atto594, 10 μM Rimeporide (Federline no. 332-12602-2), 20μM EIPA (Millipore Sigma, A3085) or 25 μM Pitstop2 (Sigma, SML 1169) mimicking our experimental setups. Subsequently, the MTT assay (Roche no. 11465007001) was performed. In brief, 10 μL of 3 - [4,5-dimethylthiazol-2-yl]-2,5-diphenyl tetrazolium bromide (MTT, 5 mg/mL) in phenol red-free neurobasal media was added to each well. The plates were incubated for an additional 4 h at 37 °C. After incubation, formazan crystals were solubilized in the provided solubilization buffer overnight at 37 °C, and the absorbance was measured at 550 nm using a Synergy H1 Multimode Microplate Reader (BioTek Instruments).

### Immunostaining and BODIPY^TM^ 493/503

Control cortical astrocytes receiving Red-C12 from CBE-treated neurons were were fixed in warm 4% PFA for 25 min at room temperature, washed twice in PBS with 0.1% Triton X-100 and blocked with PBS containing 2% bovine serum albumin and 0.2% saponin for 1 hour at room temperature. Cells were incubated with primary antibodies: anti-rabbit Plin2 (1:1,000; Novus Biologicals, NB110-40877) diluted in blocking buffer for 1 hour at room temperature. Cells were washed three times in PBS with 0.2% saponin and incubated with secondary antibodies: donkey anti-rabbit Alexa Fluor 647 (1:1,000; Thermo Fisher no. A-31573) diluted in blocking buffer for 1 h at room temperature. When staining with BODIPY^TM^ 493/503, cells were fixed in warm 4% PFA for 25 min at room temperature, incubated with 2.5μg/mL BODIPY^TM^ 493/503 for 1 hour at room temperature and washed twice with PBS. Following antibody or BODIPY^TM^ 493/503 staining, cells were washed and incubated with DAPI (1:10,000 in PBS) for 10 min at room temperature. Samples were then mounted using DAKO fluorescence mounting medium.

To validate primary cortical astrocyte and iPSC-derived astrocyte cell cultures, cells were fixed in warm 4% PFA for 25 min at room temperature, washed twice in PBS with 0.1% Triton X-100 and blocked with PBS containing 2% bovine serum albumin and 0.2% Triton X-100 for 1 h at room temperature. Cells were incubated with primary antibodies: anti-GFAP (1:1,000; Abcam, no. ab4674) and anti-Iba1 (1:1,000; Fujifilm Irvine Scientific, no. 019-19741) diluted in blocking buffer and incubated 1 hour at room temperature. Cells were washed three times in PBS with 0.1% Triton X-100 and incubated with secondary antibodies: goat anti-chicken Alexa Fluor 647 (1:1,000; Thermo Fisher no. A-21449) and goat anti-rabbit Alexa Fluor 488 (1:1,000; Thermo Fisher no. A-11008) diluted in blocking buffer for 1 h at room temperature. After washing, the cells were incubated with DAPI diluted in PBS for 10 min and mounted using DAKO fluorescence mounting media.

### GCase activity

A glucosylceramidase activity assay kit (Abcam no. ab273339) was used as per manufacturer’s instructions. Fluorescence was measured (excitation/emission = 360/445 nm) at 37 °C in endpoint mode using Synergy H1 Multimode Microplate Reader (BioTek Instruments). Three technical replicates were performed for each experiment, and all measurements were corrected by background subtraction. The activity of GCase in each sample was calculated from a standard curve and normalized to total protein levels. Experimental replicates were normalized to control.

### Imaging acquisitions and analysis

An LSM900 with Airyscan2 and and ZEN software (Zeiss) were used to acquire all images. Imaging was done with the investigator blinded to the treatments. Images related to the Red-C12 transfer, mVenus-CAAX and PFF-Atto594 transfer and LEVs experiment were acquired using plan-apochromat 40x/1.30 oil objective (Zeiss, NA = 1.4) in SR-4Y mode. Images related to Red-C12 accumulation inside cortical and dopaminergic neurons are acquired using plan-apochromat 40x/1.3 oil objective (Zeiss, NA = 1.4) in SR-2Y mode. Images related to Plin2 and BODIPYTM 493/503 experiment were acquired using plan-apochromat 63x/1.40 oil objective (Zeiss, NA = 1.4) in SR-2Y mode. Images related to Lysotracker, transferrin and Dextran were acquired using plan-apochromat 40x/1.3 oil objective (Zeiss, NA = 1.4) in SR-4Y mode. Four independent experiments and ten cells per experiment were imaged. Images related to validation of cortical and iPSC-derived astrocyte cultures and GFAP staining were acquired using plan-apochromat 20x/0.8 air objective (Zeiss, NA = 0.8). Four independent experiments and two tiles per coverslip per experiment (each covering an area of 677.24μm x 686.39μm) per experiment were imaged. Tiled images were stitched with 10% overlap. All images were Airyscan processed.

All images were analysed using the Fiji-ImageJ2 (version 2.14.0) software. The number of ectosomes containing RedC12 puncta at the soma and along the neurites were tracked through the different z-stacks and manually counted. The number, size, area and mean fluorescence intensity of Red-C12-, lysotracker-, Dextran Texas Red 70 kDa-, mVenus-CAAX- and PFF-Atto594-positive puncta in astrocytes were quantified following a maximum-intensity z-projection. Images were background substracted using a rolling-ball radius of 50 pixels, threshold using RenyiEntropy dark algorithm and analysed using the Analyze Particles function. For the mVenus-CAAX and PFF-Atto594 analyses channels were separated before applying z-projection and subsequent image processing. The mean fluorescence intensity of LysoTracker and Transferrin-647 was quantified from maximum Z-projections by drawing a region around the cell soma and measuring the intensity in that area.

Colocalization between Red-C12 puncta and either BODIPY 493/503 or Plin2 was assessed on the central optical section, defined as the plane in which the nucleus was bisected in the middle. Images were background-subtracted using a rolling-ball radius of 50 pixels and thresholded using the RenyiEntropy Dark algorithm. ROIs corresponding to Red-C12, BODIPY 493/503, and Plin2 signals were generated from the thresholded images. The overlapping area between Red-C12 and BODIPY 493/503 or Plin2 was determined using the ImageJ AND operation. The area of overlap was normalized to the total Red-C12-positive area and expressed as a percentage.

The proportions of GFAP-positive and Iba1-positive cells in astrocyte cultures were quantified from tiled images. GFAP and Iba1 signals were independently masked and used to determine the number of positive cells. The number of GFAP-positive or Iba1-positive cells was then normalized to the total number of DAPI-positive nuclei within each tile scan and expressed as a percentage.

### Statistical analyses

Statistical analysis and graphing were performed using GraphPad Prism 10. No data points were excluded from analyses. Researchers were blinded during analyses as indicated above. Data is presented as mean ± SEM, unless otherwise stated. Statistical analysis was performed on the experimental averages. To assess statistical significance, two-tailed unpaired t-test, One-way ANOVA with Šídák’s multiple comparisons test or uncorrected Fisher’s LSD were performed, as indicated in the figure legends.

## ACKNOWLEDGEMENTS

We thank Dr. Anastassia Voronova for the use of PAX6 and SOX2 antibodies for NPC astrocyte validation. Experiments were performed at the Faculty of Medicine & Dentistry Cell Imaging Core RRID: SCR_019200. MSI was supported by the Canadian Institutes of Health Research (#173321 and #191990), the Canada Research Chairs Program (#2021-00027) and a Future Leaders in Canadian Brain Research from Brain Canada (#5946). JJ was supported with a Fellowship of the Parkinson Association Alberta. TMD was supported through funding from the GBA1 Canada initiative.

## AUTHOR CONTRIBUTIONS

Conceptualization: MSI, JJ. Methodology: MSI, JJ, VS, TMD. Investigation: JJ. Technical assistance: JC, EI, DR. Visualization: MSI, JJ. Funding acquisition: MSI, JJ, TMD. Supervision: MSI, TMD. Writing – Original draft: MSI, JJ.

---

**Supplementary Figure 1.**
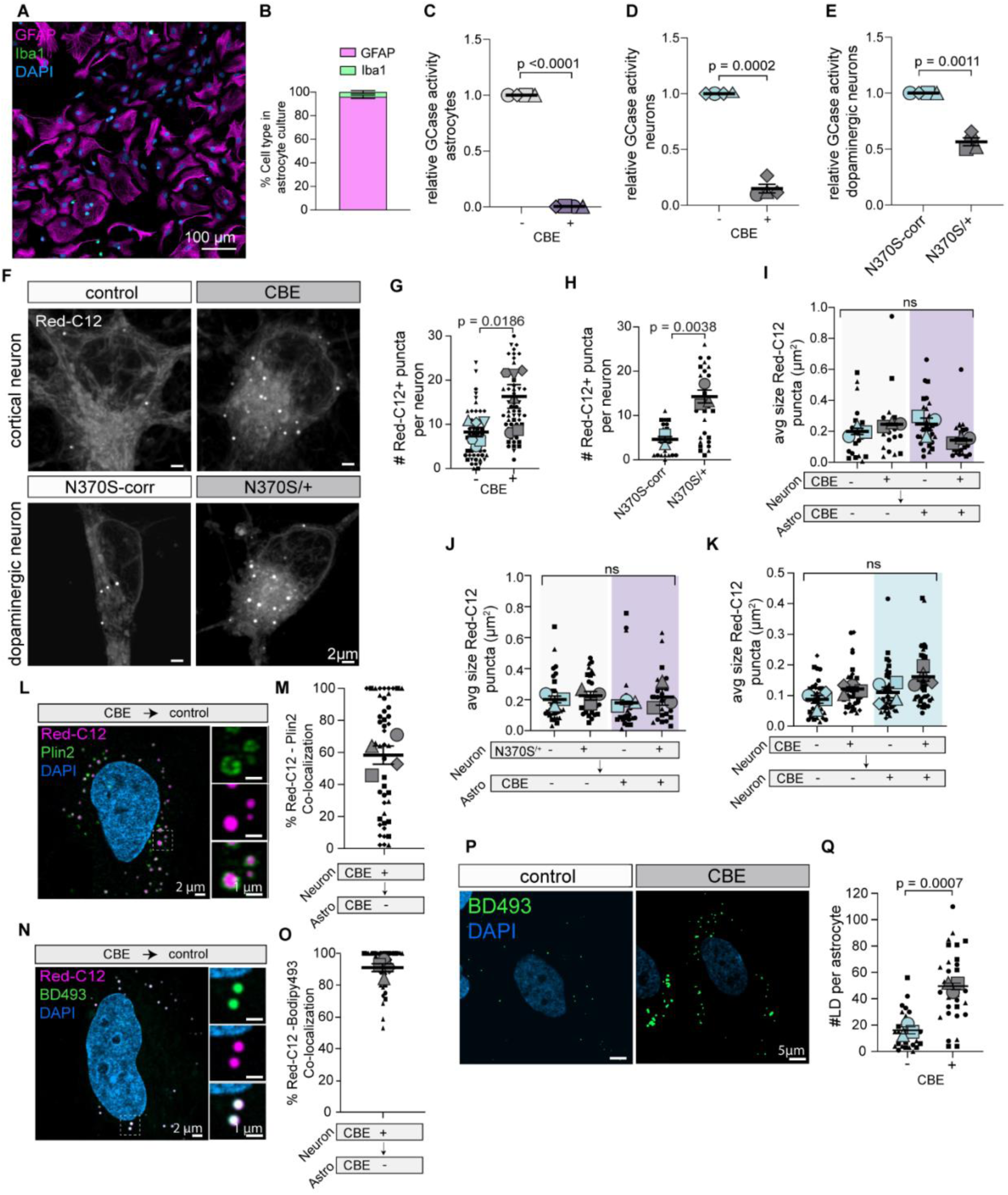
Astrocyte-specific reduction of Red-C12 uptake and accumulation in GCase deficiency. A-B. Representative tile scan of cortical astrocyte cultures with quantification of GFAP-positive astrocytes and Iba1-positive microglia. n = 3 independent experiments; 10 cells per experiment; mean ± SEM. C-E. GCase activity of cortical astrocytes and neurons treated with CBE and dopaminergic neurons normalized to their respective controls n = 4 independent experiments or inductions; mean +/- SEM.; one sample t-test. F-H. Maximum z-projections of control and GCase-deficient cortical and control or GBA1-N370S/+ dopaminergic neurons loaded with Red-C12. n=4 independent experiments; 10 cells per experiment; mean ± SEM. Unpaired *t*-test. I. Red-C12 transfer from control or CBE-treated neurons to control or CBE-treated astrocytes. Quantification of puncta size in astrocytes. n = 4 independent experiments; 10 cells per experiment; mean ± SEM. One-way Anova with Šídák’s multiple comparisons test. J. Red-C12 transfer from control or GBA1-N370S/+ neurons to control or CBE-treated astrocytes. Quantification of puncta size in astrocytes. n = 4 independent experiments; 10 cells per experiment; mean ± SEM. One-way Anova with Šídák’s multiple comparisons test. K. Red-C12 transfer from control or CBE-treated neurons to control or CBE-treated neurons. Quantification of puncta size in neurons. n = 4 independent experiments; 10 cells per experiment; mean ± SEM. One-way Anova with Šídák’s multiple comparisons test. L-M. Red-C12 transfer from CBE-treated neurons to control astrocytes. Representative image of astrocytes showing colocalization of Red-C12 puncta with Plin2. n = 4 independent experiments; 10 cells per experiment; N-O. Red-C12 transfer from CBE-treated neurons to control astrocytes. Representative image of astrocytes showing colocalization of Red-C12 puncta with BODIPY 493/503 (BD493). n = 4 independent experiments; 10 cells per experiment; P-Q. Representative images of control or CBE-treated astrocytes stained with BD493 and quantification of number of BD493-positive lipid droplets (LD) per astrocytes. n = 4 independent experiments; 10 cells per experiment; mean ± SEM. Unpaired *t*-test. Graphs are depicted as superplots where biological replicates are shown in large shapes, and technical replicates are shown as small shapes.

**Supplementary Figure 2.**
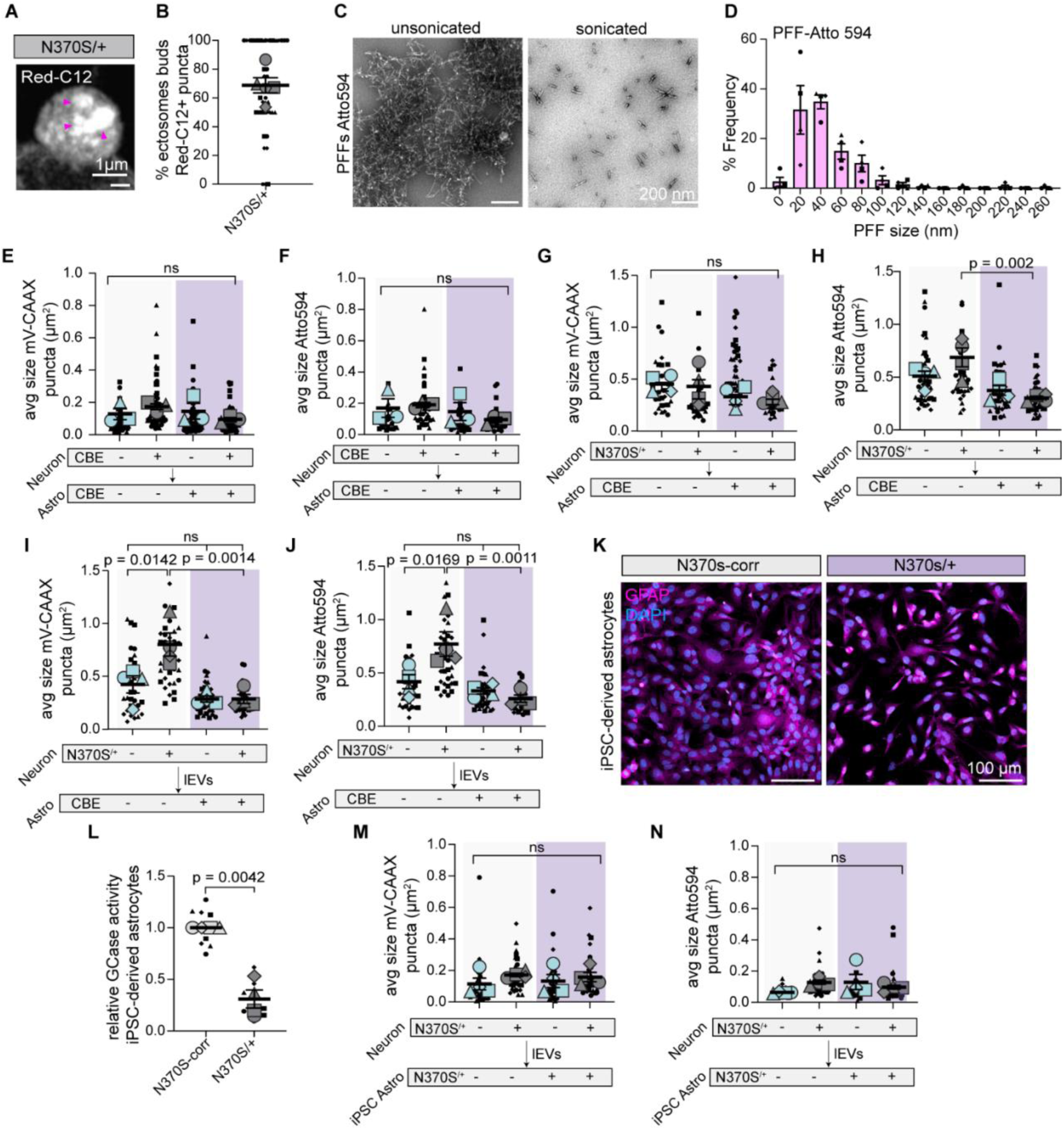
GCase deficiency in astrocytes disrupts neuron-to-astrocyte α-synuclein transport. A-B. Representative image of ectosome containing Red-C12 budding from GBA1-N370S/+ neuron. n = 4 independent experiments; 10 cells per experiment; mean ± SEM. C-D. Transmission electron micrographs of α-synuclein PFFs-Atto594 before and after sonication and the length distribution after sonication. mean ± SD. E-F. mVenus-CAAX and PFF-ATTO594 transfer from control or CBE-treated neurons to control or CBE-treated astrocytes. Quantification of puncta size in astrocytes. n = 4 independent experiments; 10 cells per experiment; mean ± SEM. One-way Anova with Šídák’s multiple comparisons test. G-H. mVenus-CAAX and PFF-ATTO594 transfer from control or GBA1-N370S/+ neurons to control or CBE-treated astrocytes. Quantification of puncta size in astrocytes. n = 4 independent experiments; 10 cells per experiment; mean ± SEM. One-way Anova with Šídák’s multiple comparisons test. I-J. Control or CBE-treated astrocytes treated with mVenus-CAAX and PFF-ATTO594 containing ectosome fractions from control or GBA1-N370S/+ neurons. Quantification of puncta size in astrocytes. n = 4 independent experiments; 10 cells per experiment; mean ± SEM. One-way Anova with Šídák’s multiple comparisons test. K. Representative tile scan of iPSC-derived astrocytes of GBA1-N370S/+ and its isogenic control stained for GFAP. L. GCase activity of iPSC-derived astrocytes with N370S/+ mutation in *GBA1* normalized to their respective controls n = 4 independent experiments or inductions; mean +/- SEM.; one sample t-test. M-N. Control or N370S/+ astrocytes treated with mVenus-CAAX and PFF-ATTO594 containing ectosome fractions from control or GBA1-N370S/+ neurons. Quantification of puncta size in astrocytes. n = 4 independent experiments; 10 cells per experiment; mean ± SEM. One-way Anova with Šídák’s multiple comparisons test. Graphs are depicted as superplots where biological replicates are shown in large shapes, and technical replicates are shown as small shapes.

**Supplementary Figure 3.**
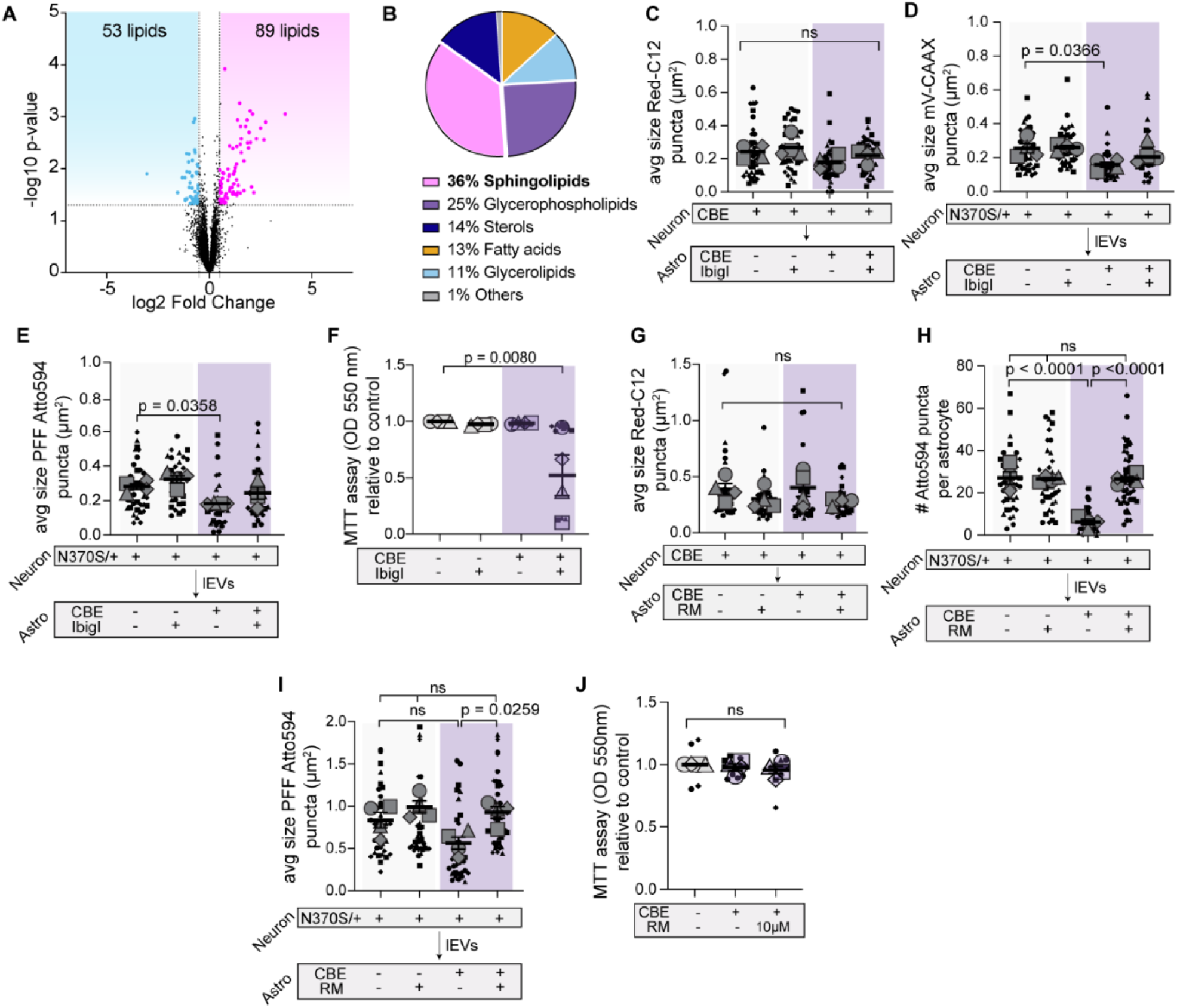
Blocking glucosylceramide synthesis or improving lysosomal function restores ectosomal α-synuclein uptake by astrocytes. A. Volcano-plot showing altered lipid species [fold change (FC) > 1.40 or < 0.71 and p < 0.05] upon CBE treatment in cortical astrocytes. B. Pie chart of lipid classes that are affected upon CBE treatment in astrocytes. C. Red-C12 transfer from CBE-treated neurons to control CBE, and/or ibiglustat-treated astrocytes. Quantification of puncta size in astrocytes. n = 4 independent experiments; 10 cells per experiment; mean ± SEM. One-way Anova with Šídák’s multiple comparisons test. D-E. Control, CBE, and/or ibiglustat-treated astrocytes treated with mVenus-CAAX and PFF-ATTO594 containing ectosome fractions from control or GBA1-N370S/+ neurons. Quantification of puncta size in astrocytes. n = 4 independent experiments; 10 cells per experiment; mean ± SEM. One-way Anova with Šídák’s multiple comparisons test. F. Cytotoxicity of astrocytes treated with CBE and/or ibiglustat by MTT assay. n = 4 independent experiments; 3 technical replicates per experiment; normalized to control; mean ± SEM; One sample *t*-test. G. Red-C12 transfer from CBE-treated neurons to control CBE, and/or rimeporide-treated astrocytes. Quantification of puncta size in astrocytes. n = 4 independent experiments; 10 cells per experiment; mean ± SEM. One-way Anova with Šídák’s multiple comparisons test. H-I. Control, CBE, and/or rimeporide-treated astrocytes treated with mVenus-CAAX and PFF-ATTO594 containing ectosome fractions from control or GBA1-N370S/+ neurons. Quantification of puncta size in astrocytes. n = 4 independent experiments; 10 cells per experiment; mean ± SEM. One-way Anova with Šídák’s multiple comparisons test. J. Cytotoxicity of astrocytes treated with CBE and/or rimeporide by MTT assay. n = 4 independent experiments; 3 technical replicates per experiment; normalized to control; mean ± SEM; One sample *t*-test. Graphs are depicted as superplots where biological replicates are shown in large shapes, and technical replicates are shown as small shapes.

**Supplementary Figure 4.**
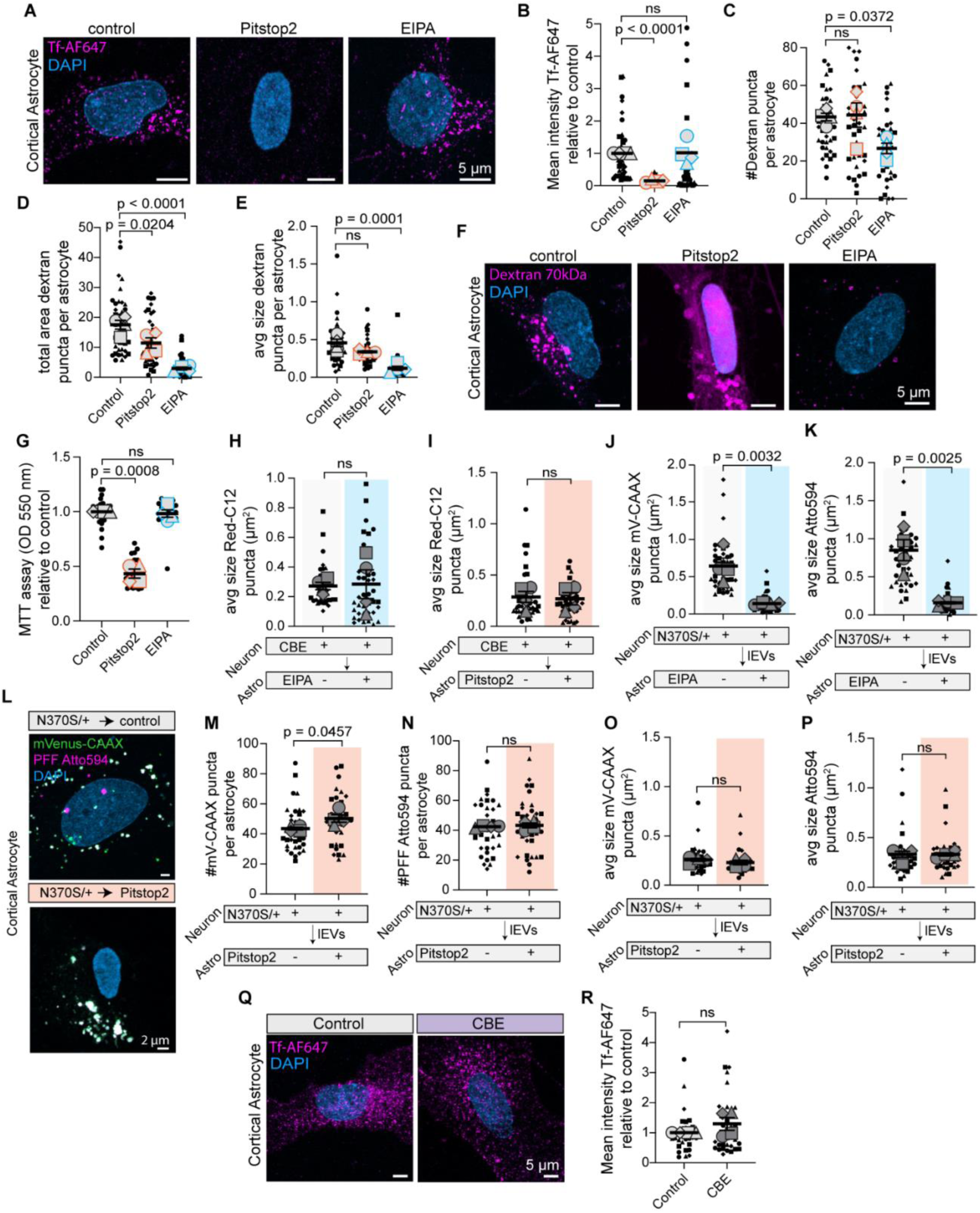
Blocking macropinocytosis but not endocytosis mimics the uptake phenotype observed in GCase deficient astrocytes. A-B. Control, EIPA, or Pitstop2 treated astrocytes were incubated with transferrin-Alexafluor647 for 1 hour. Maximum intensity projection of representative image. Mean intensity of Transferrin normalized to total area of astrocyte. n = 4 independent experiments; 10 cells per experiment; mean ± SEM. One-way Anova with Šídák’s multiple comparisons test. C-F. Control, EIPA, or Pitstop2 treated astrocytes were incubated with Dextran 70kDa for 1 hour. Maximum intensity projection of representative image. Number, total area and size of Dextran puncta per astrocyte were quantified. n = 4 independent experiments; 10 cells per experiment; mean ± SEM. One-way Anova with Šídák’s multiple comparisons test. G. Cytotoxicity of astrocytes treated with Pitstop2 or EIPA by MTT assay. n = 4 independent experiments; 3 technical replicates per experiment; normalized to control; mean ± SEM; One sample *t*-test. H. Red-C12 transfer from CBE-treated neurons to control or EIPA-treated astrocytes. Quantification of puncta size in astrocytes. n = 4 independent experiments; 10 cells per experiment; mean ± SEM. Unpaired *t*-test. I. Red-C12 transfer from GBA1-N370S/+ neurons to control or Pitstop2-treated astrocytes. Quantification of puncta size in astrocytes. n = 4 independent experiments; 10 cells per experiment; mean ± SEM. Unpaired *t*-test. J-K. Control, or EIPA-treated astrocytes treated with mVenus-CAAX and PFF-ATTO594 containing ectosome fractions from control or GBA1-N370S/+ neurons. Quantification of puncta size in astrocytes. n = 4 independent experiments; 10 cells per experiment; mean ± SEM. Unpaired *t*-test. L-P. Control, or Pitstop2-treated astrocytes treated with mVenus-CAAX and PFF-ATTO594 containing ectosome fractions from control or GBA1-N370S/+ neurons. Representative maximum intensity projection. Quantification of puncta number and size in astrocytes. n = 4 independent experiments; 10 cells per experiment; mean ± SEM. Unpaired *t*-test. Q-R. Control, or CBE-treated astrocytes were incubated with transferrin-Alexafluor647 for 1 hour. Maximum intensity projection of representative image. Mean intensity of Transferrin normalized to total area of astrocyte. n = 4 independent experiments; 10 cells per experiment; mean ± SEM. Unpaired *t*-test. Graphs are depicted as superplots where biological replicates are shown in large shapes, and technical replicates are shown as small shapes.

